# Hierarchical Brain–LLM Alignment Reveals Layer-Specific Neural Representations of Second Language Proficiency

**DOI:** 10.1101/2025.06.17.660057

**Authors:** Rieko Kubo, Shinji Nishimoto, Tomoya Nakai

**Affiliations:** Center for Information and Neural Networks, National Institute of Information and Communications Technology, Suita, Japan; Graduate School of Medical and Dental Sciences, Institute of Science Tokyo, Tokyo, Japan; Graduate School of Frontier Biosciences, The University of Osaka, Suita, Japan; Graduate School of Medicine, The University of Osaka, Suita, Japan; Araya Inc., Tokyo, Japan

**Author notes:** Corresponding author, Full postal address: 6F Sanpo Sakuma Building, 1-11 Kandasakumacho, Chiyoda-ku, Tokyo, Japan.

**Keywords:** fMRI, encoding model, individual variability, second language (L2), language model

## Abstract

Second language (L2) comprehension is thought to proceed hierarchically, from basic word recognition to complex discourse-level understanding. However, the neural mechanisms underpinning this hierarchical progression remain poorly understood. Leveraging the hierarchical nature of linguistic representations in large language models (LLMs), we investigated how L2 proficiency modulates layer-wise brain-LLM alignment. Using functional magnetic resonance imaging (fMRI) data collected during discourse listening from 54 participants with varying levels of L2 proficiency, we constructed individualized encoding models to quantify how proficiency shapes the brain-LLM representational correspondence. In high-proficiency individuals, LLM-based models reliably predicted neural responses in a distributed network comprising temporal, frontal, and medial-parietal cortices, while for low-proficiency participants, prediction performance decreased. Notably, alignment in the superior temporal sulcus (STS) showed a robust correlation with L2 proficiency at mid-to-deep LLM layers, suggesting that the STS encodes higher-level linguistic abstractions essential for advanced language comprehension. These findings indicate that L2 proficiency modulates the alignment between LLMs and the brain in a graded, layer-dependent manner, providing insights into hierarchical neural language representations.

## 1. Introduction

There is considerable variability in the individual ability to understand second languages (L2), from near-native proficiency to inability. L2 proficiency assessments, such as the Common European Framework of Reference for Languages (CEFR), characterize this progression as a shift from basic phrase recognition to more integrated, discourse-level processing^1^. Complementing this hierarchical behavioral framework, recent neuroimaging studies have shown that neural responses to linguistic information vary depending on L2 proficiency^2–5^. These findings highlight a dynamic reorganization of brain activity with increasing L2 proficiency, often involving the progressive recruitment of distributed cortical networks across frontal and temporal regions.

Such hierarchical brain organization has also been described in first language (L1) processing, where low-level features are typically processed in localized brain regions, while higher-order information (e.g., narrative-level integration) recruits more distributed cortical networks^6–16^. For example, phonological and lexical processing tends to engage superior temporal gyrus (STG)^17,18^, while syntactic and semantic integration recruits additional frontal and temporal areas, such as the inferior frontal gyrus^14,19^. Furthermore, discourse-level comprehension has been associated with even broader networks, including medial prefrontal and parietal cortices^13^, suggesting a progressive expansion of neural recruitment as linguistic complexity increases. However, the neural mechanisms underlying this hierarchy, as well as the representational reorganization associated with L2 proficiency, remain poorly understood.

A major technical challenge in previous studies has been isolating intermediate stages within the L2 processing hierarchy^20^. One promising approach for overcoming this limitation involves the integration of encoding models^21,22^ with large language models (LLMs) such as GPT-2^23^. Trained on a vast corpus of real-world text, LLMs capture hierarchical linguistic representations, including syntactic structures, semantic relationships, and contextual dependencies^23–26^. Recent studies have shown that LLM-derived features can predict brain activity during naturalistic language comprehension with remarkable accuracy^27–36^, suggesting a representational alignment between LLMs and the human brain. More importantly, this quantitative methodology enables connecting brain activity to behavioral measures^30,37,38^ by examining the layer-wise relationship between prediction performance and L2 comprehension scores, thereby uncovering the neural correlates of hierarchical language processing. Despite this progress, most prior research on brain-LLM alignment has focused on L1 comprehension, largely overlooking the unique variability in L2 proficiency. Even in recent efforts to apply brain-LLM alignment to L2 comprehension^39^, the relationship between hierarchical representations and L2 proficiency has not been examined.

To address this gap, we conducted an fMRI study with 54 native Japanese speakers listening to L2 discourses (English) (**Fig. 1**). Using voxel-wise encoding models^21^, we examined how the brain-LLM alignment varies across LLM layers and how this alignment is modulated by L2 proficiency. The encoding models incorporated latent features extracted from different layers of GPT-2^23^, an autoregressive LLM widely used in brain-LLM alignment studies^30–32^. We then quantitatively linked the prediction accuracy of these models to L2 proficiency. We hypothesized that individuals with lower L2 proficiency would exhibit a stronger alignment with shallow LLM layers, while those with higher proficiency would show greater alignment with middle to deeper layers. Furthermore, we expected a pronounced proficiency-dependent alignment in brain regions previously implicated in hierarchical language processing. Finally, leveraging the flexibility of encoding models—a key advantage of the approach—, additional analyses using LLMs with different training objectives and representational characteristics from GPT-2 were conducted to examine whether distinct architectural designs yield different layer-wise alignment with L2 proficiency. Together, the current findings advance our understanding of how L2 proficiency shapes the brain’s hierarchical language architecture.

**Fig. 1.**
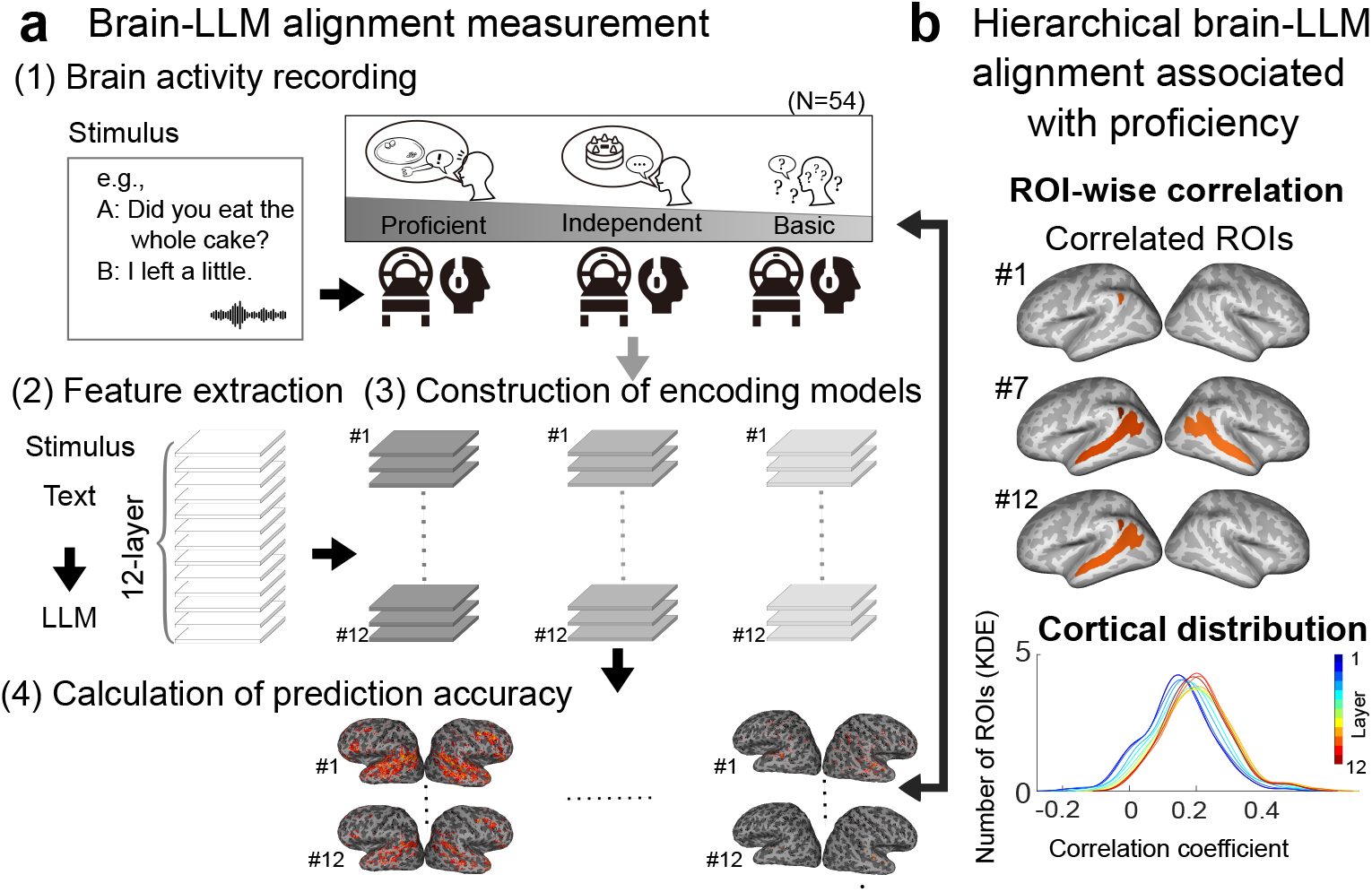
Study overview. Schematic representation of the experimental workflow. **a** Measurement of brain-LLM alignment. (1) The neural activity of participants with varying L2 proficiency was recorded using fMRI while they engaged in L2 discourse comprehension tasks. (2) Latent features of the stimuli were extracted from each LLM layer (primarily 12-layer GPT-2). (3) Voxel-wise encoding models were constructed for each layer and participant to map LLM-derived features onto brain activity, enabling a detailed layer-by-layer analysis. (4) Encoding model performance was calculated at each layer for each participant. **b** Hierarchical brain-LLM alignment as a function of L2 proficiency. Encoding model performance within each region of interest (ROI) was correlated with L2 proficiency to identify localized brain regions and model layers showing proficiency-related modulation. To capture global patterns, the cortical distribution of these correlations was examined using a kernel density estimation.

## 2. Results

### 2.1. Both L2 proficiency and LLM layers affect brain-LLM alignment

We first examined whether the predictability of brain activity by LLM features is affected by individual language proficiency. To this aim, we divided the participants into three proficiency levels (Proficient users, N = 20; Independent users, N = 26; Basic users, N = 8) based on independent English test scores (see section **4.2**.). **Fig. 2** illustrates the impact of GPT-2 layers and L2 proficiency levels on prediction performance. The proportion of significant voxels varied across L2 proficiency levels and GPT-2 layers (**Fig. 2a**), suggesting that both factors influence model alignment with brain activity during L2 processing. GPT-2 models predicted a large portion of language-related regions for Proficient users, and smaller regions for Independent and Basic users (mean percent of significant voxels across the cortex and layers, Proficient users, 16.7%; Independent users, 12.8%; Basic users, 9.1%).

**Fig. 2.**
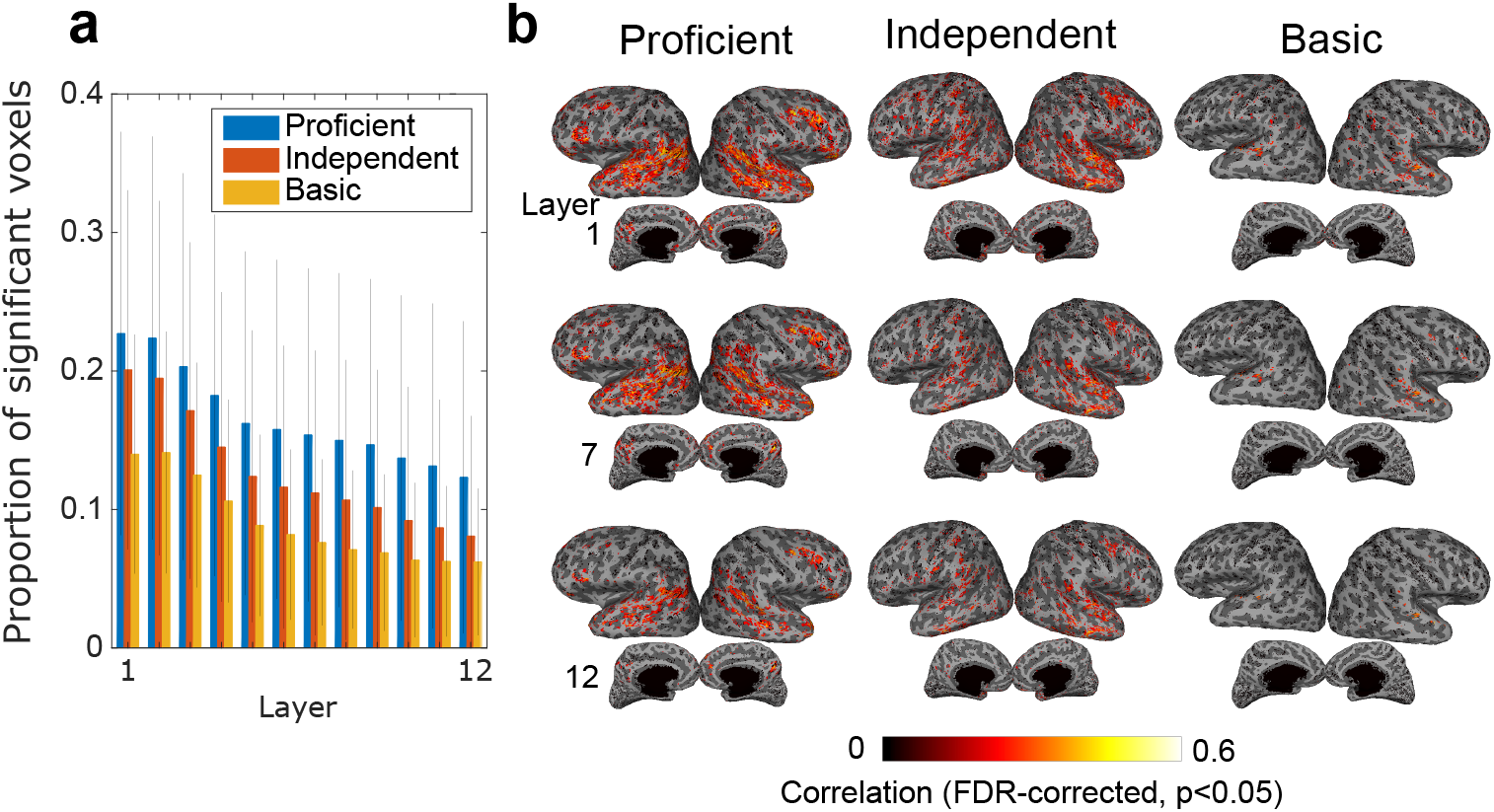
Effect of language proficiency and GPT-2 layers on prediction performance. **a** Proportions of significant voxels across 12 GPT-2 layers and three L2 proficiency levels (Proficient, Independent, Basic). Error bar, SD. **b** Example cortical maps of prediction accuracy for a representative participant from each of the three proficiency levels and three layers. Only significant voxels are shown (*p* < 0.05, false discovery rate (FDR)-corrected). Maps depict inflated cortical surfaces with the left and right hemispheres displayed on the corresponding sides, respectively. Upper and lower rows show lateral and medial views, respectively.

To complement these group-level findings, we also visualized individual brain maps of prediction accuracy to examine their alignment across the cortex. The prediction accuracy of different GPT-2 layers (layers 1, 7, and 12) in representative Proficient, Independent, and Basic users demonstrate distinct patterns of brain-LLM alignment across a wide range of regions (**Fig. 2b**). Proficient participants exhibited larger and more widespread alignment, encompassing not only areas traditionally implicated in language processing, such as the superior temporal sulcus (STS), and middle temporal gyrus (MTG), but also regions associated with broader semantic and integrative functions, including the angular gyrus (AG) and precuneus (PCu). In contrast, participants with lower proficiency showed progressively lower engagement in these regions. These findings suggest a hierarchical and distributed nature of L2 processing as well as its modulation by language proficiency.

### 2.2. Layer-specific prediction patterns are modulated by L2 proficiency

To further investigate how L2 proficiency influences the language representation hierarchy, we identified the best-fitting LLM layer for each voxel and calculated the proportion of voxels corresponding to each layer within predefined regions of interest (ROIs) (see section **4.9**.). Example whole-cortex maps illustrate distinct layer fitting patterns across the three proficiency groups (**Fig. 3a**). Broader brain regions were predicted by deeper layer features in Proficient and Independent users compared to Basic users. In addition, ROI-based analysis provided the proportion of voxels predicted by each encoding model. We focused on language-related ROIs previously associated with L1 comprehension (STS, supramarginal gyrus [SMG], AG, MTG, inferior frontal gyrus triangularis [IFGtri], and PCu)^30^ (**Fig. 3b**). Across all these ROIs, we found patterns consistent with the whole-brain analysis: compared to Independent and Basic users, Proficient users had a high proportion of voxels best predicted by deeper layers.

**Fig. 3.**
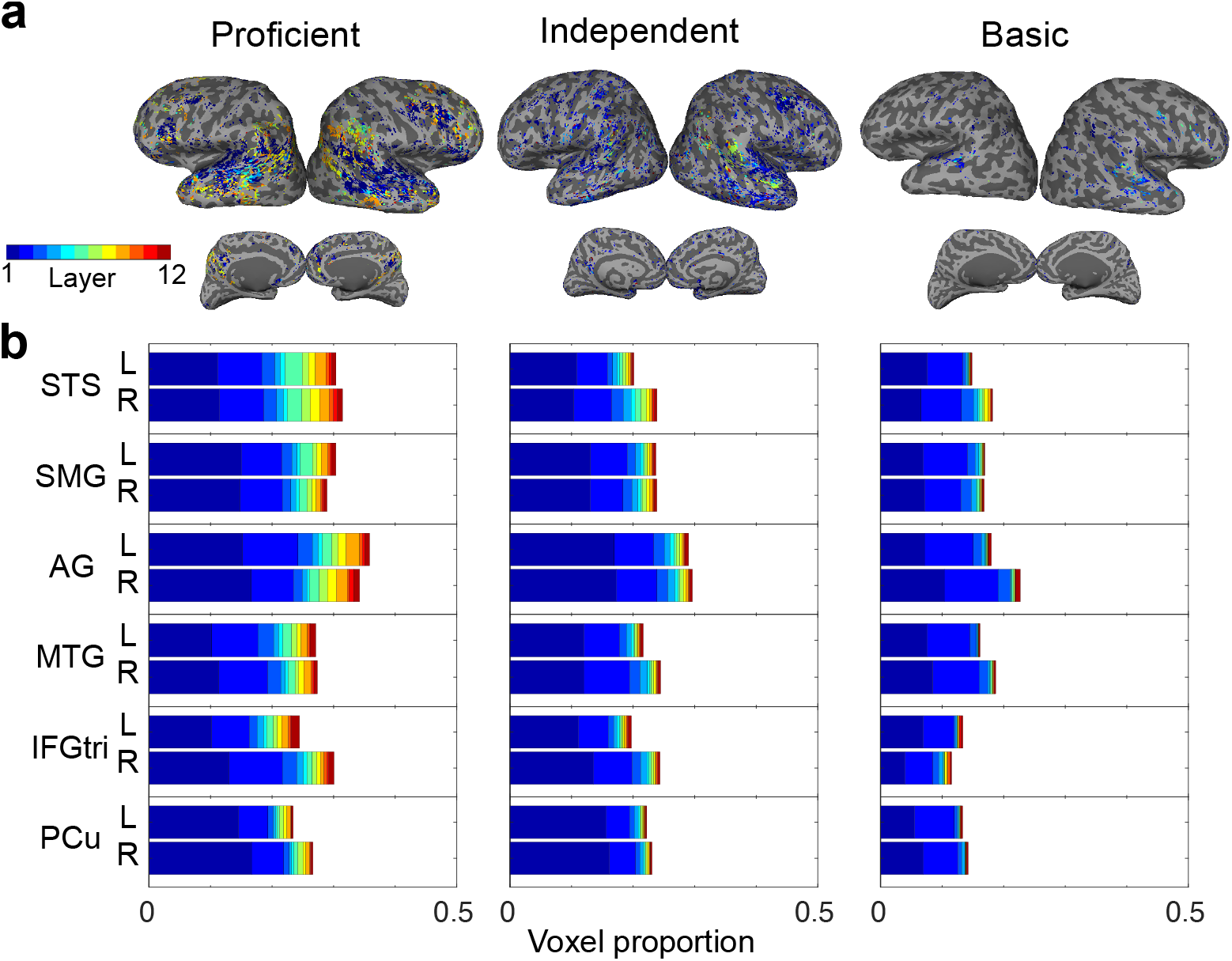
Best-fitting model for each voxel across GPT-2 layers. a Example brain maps showing the best-fitting GPT-2 encoding model (out of 12 layers) for each voxel in a representative participant from each proficiency level (Proficient, Independent, Basic). The color gradient represents the corresponding best-fit layer. b Mean proportion of voxels assigned to the best-fitting model within specific regions of interest (ROIs) in each hemisphere for different L2 proficiency levels. STS, superior temporal sulcus; SMG, supramarginal gyrus; AG, angular gyrus; MTG, middle temporal gyrus; IFGtri, inferior frontal gyrus triangularis; PCu, precuneus, L; left hemisphere, R, right hemisphere.

### 2.3. Prediction accuracy correlates with L2 proficiency in deeper LLM layers

To further examine the impact of L2 proficiency in each region, we calculated the correlation between prediction accuracy and L2 proficiency scores across GPT-2 layers and anatomical ROIs (see section **4.10**.). We first identified anatomical ROIs for which L2 proficiency was significantly correlated with prediction accuracy in specific GPT-2 layers (**Fig. 4a**; shown for four representative layers: layers 1, 5, 9, and 12, results across all layers are shown in **Supplementary Fig. S1a**). Significant correlations were observed in regions associated with the integration of contextual and social information^40–43^, including the bilateral STS, Jensen sulcus, and subparietal areas. Notably, in these ROIs, more extensive correlation patterns were observed in deeper layers. These findings suggest that the GPT-2 model captures neural correlates of L2 proficiency that reflect hierarchical linguistic representations^44,45^.

**Fig. 4.**
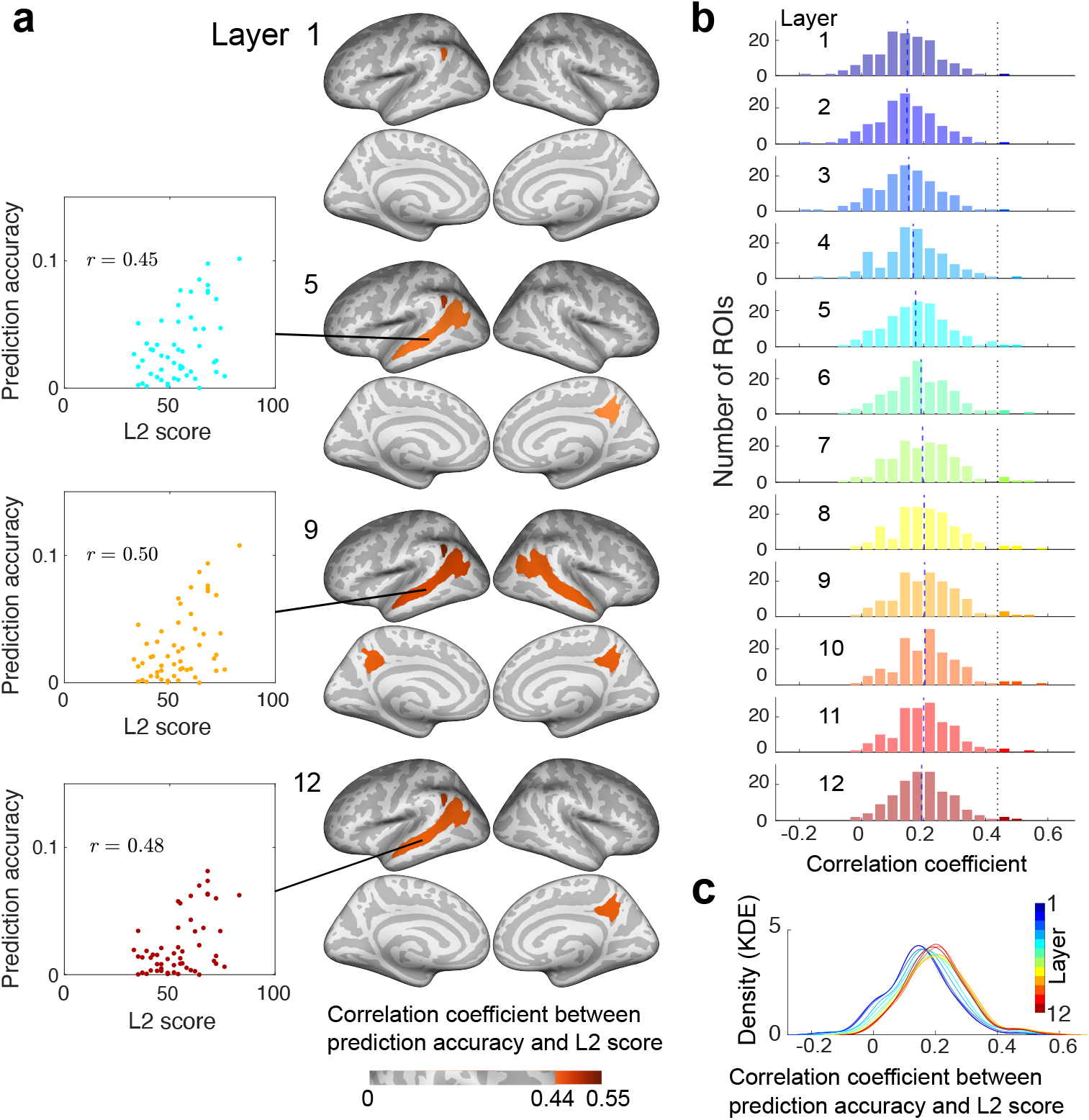
Correlation between prediction accuracy and L2 proficiency. **a** Brain map showing ROIs with significant correlations between L2 proficiency and prediction accuracy (color-coded, *p* < 0.05, FDR corrected), accompanied by illustrative scatter plots. **b** Distribution of correlation coefficients between L2 proficiency and prediction accuracy across the 12 GPT-2 layers. The black dotted line indicates the significance threshold (*r* = 0.44), and the blue dashed line the median for each layer. **c** Kernel density estimation of the distribution of correlation coefficients across all layers.

Moreover, the distribution of correlation coefficients demonstrates how language proficiency modulates hierarchical representations across all cortical ROIs (**Fig. 4b**). Shallower layers (e.g., layers 1–3) exhibited a lower range of correlation coefficients (median = 0.15, 0.15, 0.15, respectively), suggesting limited and less variable associations across brain regions. In contrast, deeper layers (e.g., layers 10–12) showed a relatively higher distribution centered around a moderate positive correlation (median = 0.20, 0.20, 0.19, respectively). Kernel density estimation further confirmed this trend, demonstrating an upward shift of the distribution peak (**Fig. 4c**). These findings indicate a stronger association between LLM-brain alignment and L2 proficiency in deeper layers.

### 2.4. Distinct and shared correlation patterns across LLMs

To investigate whether different LLMs exhibit distinctive patterns explaining L2-related brain activity, we tested two other LLMs beyond GPT-2 (**Fig. 5**). Specifically, we used XLNet^46^ and Llama 2^47^, both sharing Transformer architecture with GPT-2. First, we calculated layer-wise correlations between prediction accuracy and L2 proficiency (**Fig. 5a** shows results for a mid-to-deep layer, defined as 0.75 of the total layer depth (e.g., layer 9 for a 12-layer model, or layer 24 for a 32-layer model), results across all layers are shown in **Supplementary Figs. S1b, c**) and obtained distinct correlation profiles across LLMs.

**Fig. 5.**
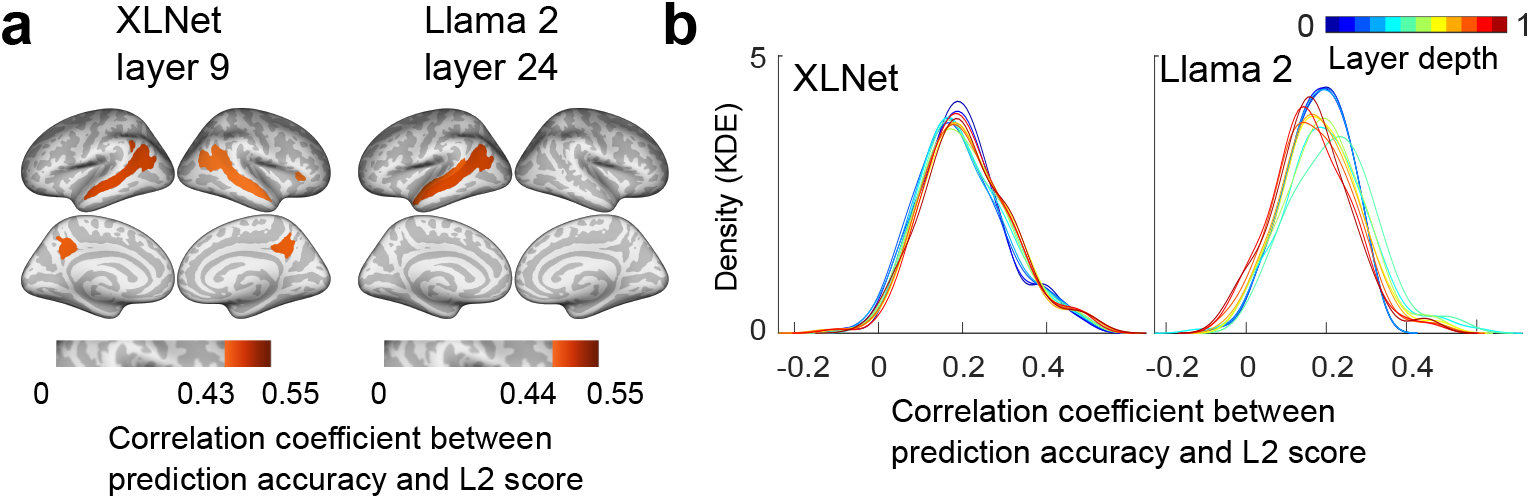
Correlation between prediction accuracy and L2 proficiency across LLMs. **a** Brain map showing ROIs with significant correlations between L2 proficiency and prediction accuracy (color-coded, *p* < 0.05, FDR corrected). A representative mid-to-deep layer is shown here; full layer-wise results are available in Supplementary Figure S1b, c. **b** Kernel density estimation (KDE) of the distribution of correlation coefficients between prediction accuracy and L2 proficiency. These KDEs represent the overall distribution of correlation coefficients obtained from all ROIs, calculated across all layers of the XLNet and Llama 2 encoding models, illustrating the layer-wise patterns of alignment with L2 proficiency.

In contrast, we observed substantial overlap in the sets of correlated ROIs across all models, including the STG, Jensen sulcus, STS, subparietal sulcus, and the anterior horizontal fissure (**Fig. 5a**). These findings suggest that, although model architecture influences the specific patterns of brain-LLM alignment, certain neural representations of language processing are robustly captured across different Transformer-based LLMs.

## 3. Discussion

In the present study, we investigated the hierarchical mechanisms underlying L2 proficiency and its neural representations. To this end, we constructed a series of voxel-wise encoding models incorporating latent features extracted from different layers of GPT-2, an autoregressive LLM widely used in brain-LLM research. This approach enabled us to quantitatively link neural activity during L2 comprehension to LLM-derived features, and to examine how individual variability in L2 proficiency influences hierarchical language representations in the brain.

Our findings revealed that L2 proficiency modulates the brain-LLM alignment (**Figs. 2, 3**). In Proficient users, this alignment extended beyond the temporal cortex to broader regions of the parietal cortex, including the inferior parietal lobule and superior parietal lobule^9,48,49^, and was more strongly aligned with deeper GPT-2 layers. In contrast, users with lower proficiency exhibited a weaker alignment, primarily confined to the temporal lobes and more prominent in shallower layers. With increasing L2 proficiency, the neural representations became more distributed across cortical regions, potentially facilitating more complex semantic and contextual comprehension. These patterns are consistent with previous studies in native language processing^6,50–53^. Further, these results support the hypotheses that deeper LLM layers capture complex linguistic features^23–26^, and that individual differences in L2 proficiency modulate such hierarchical neural representations^54,55^.

In addition, we showed that L2 proficiency correlates with brain-LLM alignment in both localized and distributed brain regions. The impact of L2 proficiency was particularly evident in the STS, Jensen sulcus, and subparietal areas, which are implicated to be involved in word-level comprehension, information integration, and contextual and social processing (**Fig. 4a**)^9,40,56,57^. In shallower LLM layers, a brain-LLM alignment associated with L2 proficiency was observed in the left STS, while intermediate layers exhibited alignment in the bilateral STS, suggesting a hierarchical propagation of linguistic information^58–62^. Interestingly, while the STS showed a significant alignment with L2 proficiency, other language-sensitive regions such as the IFG did not exhibit a comparable pattern. This discrepancy may reflect functional differences between these regions or be due to the nature of the conversational speech stimuli used in the present study, which emphasize semantic and contextual comprehension over syntactic processing^9,40,56,59–62^. The STS is known to process broad, context-dependent information, including nonverbal intent such as social and contextual cues^9,40,56^, whereas the IFG is more closely associated with syntactic integration^63,64^. These findings may therefore reflect the relative contribution of lexical-semantic content versus syntactic structure in shaping the brain-LLM alignment^65^. Despite these ROI-specific patterns, we found a gradual increase in the association between brain-LLM alignment and L2 proficiency across the cortex (**Fig. 4b**), suggesting that language understanding is supported by distributed cortical processes and representational hierarchies.

Furthermore, a comparative analysis across different LLMs (GPT-2, XLNet, and Llama 2) revealed distinct and common patterns in the way these models relate to L2 proficiency, as shown by the distribution of correlation coefficients and identified cortical regions (**Figs. 4, 5**). These results indicate that LLMs with different architectures have distinct representational characteristics. As shown by the kernel density estimations in **Figs. 4c, 5b**, the distribution of correlation coefficients across layers reveals varying patterns: one model shows a shift toward higher correlations in deeper layers, while others exhibit a more uniform or mid-layer peak in correlation strength. This illustrates model-specific differences in how linguistic features relevant to L2 proficiency are processed and represented across their internal hierarchies. Leveraging such architectural differences could provide insights into how specific aspects of language processing are modulated by L2 proficiency. Despite these inter-model differences, the overlapping ROIs across all models suggest consistent capture of certain neural representations regardless of the LLM used. This consistency may reflect shared underlying mechanisms of language feature encoding in both artificial models and the human brain.

The observed brain-LLM alignment can be interpreted through computational frameworks such as generalizable representation^66^ and predictive coding^27,32,67,68^. The generalizable representations framework^66^ posits that intermediate LLM layers capture versatile features not specialized in prediction, leading to alignment between the brain and the LLM. Accordingly, the brain-LLM alignment would show the strongest association with L2 proficiency at intermediate layers. However, our findings revealed no such peak at middle layers (**Figs. 4b, c**), suggesting that the brain-LLM alignment is not solely driven by versatile feature extraction. Rather, it may reflect hierarchical information refinement across successive LLM layers. This pattern was particularly evident in GPT-2 (**Figs. 4, 5**), a key model in predictive coding research^30–32^. These results underscore the relevance of gradual information refinement and predictive processing in shaping brain-LLM alignment.

Despite these insights, several limitations must be acknowledged. First, our study did not include L1 data, which limits our ability to directly compare L1 and L2 comprehension^39^ or investigate L1–L2 interactions^69^. However, previous studies have shown that bilinguals often recruit overlapping neural structures for both L1 and L2 processing^70,71^. Moreover, recent research has identified a universal language network involving shared neural substrates, regardless of linguistic background^72^. Despite possible neural differences between L1 and L2 processing^70,71^, particularly among less proficient L2 users, these findings suggest shared core neural mechanisms underlying language comprehension across L1 and L2. This supports the use of L2-based approaches to investigate the neurobiological underpinnings of language processing. Second, the lower prediction accuracy observed in low-proficiency participants may reflect lower engagement rather than differences in language processing. However, this explanation appears unlikely. Shallower layers, which are more sensitive to lower-level linguistic features, still exhibited reasonable prediction accuracy across all proficiency levels. This suggests that basic language processing is preserved regardless of L2 proficiency. In contrast, proficiency-related differences were more prominent in deeper layers, which are associated with higher-order linguistic integration. These results indicate that the observed differences are more likely attributable to variability in hierarchical language processing, rather than to general attentional factors. Finally, although our findings highlight hierarchical brain-LLM alignment, the precise linguistic functions captured by these encoding models remain unclear. While a recent study addressed this issue^73^, future research should aim to further elucidate how neural representations evolve across LLM layers and how these processes interact with L2 proficiency.

In summary, our findings indicate that L2 proficiency hierarchically modulates the brain-LLM alignment. Specifically, alignment in the STS correlates with L2 proficiency, suggesting that the STS encodes higher-level linguistic abstractions essential for advanced language comprehension. Additionally, stronger correlation in deeper LLM layers across cortical regions suggest that representational hierarchies and distributed cortical processes may support the progression from basic word recognition to complex discourse-level comprehension. Together, our study successfully linked hierarchical representations to behavioral measures of L2 proficiency, providing insights into how L2 learning shapes neural language representations.

## 4. Methods

### 4.1. Participants

Fifty-four healthy native Japanese speakers, aged 19–32 years (24 females), participated in the experiment. They were all right-handed (laterality quotient = 54–100), as assessed using the Edinburgh inventory^74^, and self-reported no prior diagnosis of language or auditory abnormalities. Written informed consent was obtained from all participants prior to their participation in the experiment. This experiment was approved by the ethics and safety committee of the National Institute of Information and Communications Technology in Osaka, Japan.

### 4.2. Assessing second-language proficiency

English proficiency was assessed using the EF Standard English Test (EF SET), a 50-min online assessment of reading and listening comprehension skills (https://www.efset.org/). The participants received individual scores on a scale of 0–100 on both reading and listening comprehension, as well as an overall score over the same range. The overall score corresponds to a proficiency level on the 3-level CEFR. The participants’ overall scores of EF SET and CEFR levels varied widely, ranging from 32 (“Basic user”) to 82 (“Proficient user”).

To further validate proficiency, apart from the behavioral data acquired during the fMRI (TOEFL listening test) and the EF SET scores from all participants, TOEIC scores obtained in the past two years were collected from 51 participants. Pearson correlations were calculated among the following three test scores: (i) comprehension score during fMRI (TOEFL questions); (ii) EF SET listening score; (iii) TOEIC listening score, excluding missing data. The correlations ranged from 0.75 to 0.76, indicating that the acquired data aligned with the participants’ general language proficiency.

In addition, we utilized the EF SET listening comprehension scores as a measure of proficiency since the study employed listening tasks during fMRI scanning. Proficiency levels (i.e., CEFR levels) were assigned based on the EF SET listening scores (0–40: Basic user [∼A2 levels], 41– 60: Independent user [B1–B2 levels], 61-100: Proficient user [C1–C2 levels]). As a result, 20 participants were classified as Proficient user, 26 as Independent user, and 8 as Basic user.

### 4.3. Stimuli and procedure

To explore daily-life comprehension, we selected stimuli from the listening comprehension section of the TOEFL iBT TESTS (Official Guide to the TOEFL Test, ISBN-13 978-1260473353, 978-1260470338). We included 15 conversations involving two speakers, each accompanied by two comprehension questions. These questions were intended to assess: 1) basic comprehension (the main idea or purpose of a conversation), 2) connecting information (identifying the organization of information to make connections between important points or inferencing based on important points), and 3) pragmatic understanding (identifying a speaker’s purpose in making a statement or their attitude, opinion, or degree of certainty). Answering those questions requires abilities in association, working memory, inference, and social cognition in addition to basic speech/language skills. For each conversation, we selected two comprehension questions.

Each fMRI run consisted of a single conversation. Fifteen runs were conducted for each participant. At the beginning of each run, 6 s of dummy scans were acquired, during which the fixation cross was displayed; these dummy scans were later omitted from the final analysis to reduce noise. Participants were asked to fixate on a fixation cross presented at the center of the screen and to listen to a conversation through MRI-compatible ear tips. Following each conversation, two comprehension questions were presented; participants were required to select the appropriate response from four choices displayed on the screen by pressing the corresponding button. Each question was displayed for 15 s. Scans obtained during the question-answering phase were later omitted from encoding model analysis. The percentage of correct answers for comprehension questions was calculated for each participant by averaging 30 questions (two questions * 15 runs). The percentage of correct answers ranged from 0.1 to 0.9 and correlated well with other English proficiency indices (see section **4.2**.).

The speech signals were presented binaurally using Sensimetrics S14 in-ear piezoelectric headphones (Sensimetrics). They were filtered to correct the headphones’ frequency response using a custom equalization filter provided by the manufacturer. Stimulus presentation and behavioral data collection were managed using presentation software from Neurobehavioral Systems (Albany, CA, USA). Optic response pads with two buttons on each of the left and right hands were used to measure button responses (HHSC-2 × 2, Current Designs, Philadelphia, PA, USA).

### 4.4. fMRI data acquisition

The experiment was conducted on a 3.0T MRI scanner (MAGNETON Prisma-Fit; Siemens, Erlangen, Germany), equipped with a 64-channel head coil. We obtained 78 2.0-mm-thick interleaved axial slices without a gap, using a T2-weighted, gradient-echo, multiband, echo-planar imaging sequence (repetition time [TR] = 1,000 ms, echo time [TE] = 30 ms, flip angle [FA] = 60º, field of view [FOV] = 192 × 192 mm^2^, voxel size = 2 × 2 × 2 mm^3^, multiband factor = 6). The number of volumes collected for each run varied depending on the stimulus length (mean volumes ± SD: 204 ± 19 s, including 6 s of initial dummy scans and 30 s of comprehension-question scans at the end of each run). Additionally, for anatomical reference, high-resolution T1-weighted images of the whole brain were acquired from all participants using a magnetization-prepared rapid acquisition gradient-echo sequence (MPRAGE, TR = 2,530 ms, TE = 3.26 ms, FA = 9°, FOV = 256 × 256 mm^2^, voxel size = 1 × 1 × 1 mm^3^).

### 4.5. fMRI data preprocessing

Motion correction was performed in each run using the statistical parametric mapping toolbox (SPM8; Wellcome Trust Center for Neuroimaging, London, UK; http://www.fil.ion.ucl.ac.uk/spm/). All volumes were aligned to the first EPI image for each participant. Low-frequency drift was removed using a median filter with a 120-s window and the response for each voxel normalized by subtracting the mean response and scaling it to the unit variance. We used FreeSurfer^75^ to identify cortical surfaces from anatomical data and to register them to functional data voxels. For each participant, the voxels identified in the cerebral cortex were used for analysis (54,288–76,333 voxels per participant). We defined 148 anatomical ROIs based on the Destrieux cortical atlas^76^.

### 4.6. Speech stimuli preprocessing

We used the Penn Phonetics Lab Forced Aligner (P2FA; Yuan2008P2FA) to automatically align the speech signals with the transcript obtained from the TOEFL official guidebook. The Carnegie Mellon University Pronouncing Dictionary (CMU Dictionary) was used for alignment at both word and phonemic levels. Subsequently, the aligned transcripts underwent manual verification and correction using Praat^77^. During manual correction, we also referenced the word- and phonemic-level output of another segmentation tool (webMAUS^78,79^). Then, the aligned transcripts were transformed into lists of words with their onset and offset times.

### 4.7. Feature extraction from LLMs

In this study, we used GPT-2^23^, a unidirectional-attention Transformer model, as primary LLM as it was previously used for testing brain-aligned language models^27,32,66^. Transcripts were tokenized using GPT-2 tokenizer and input into a pre-trained GPT-2 model provided by Hugging Face (“gpt2”), featuring 12 layers and 768 hidden units.

Given that participants contextualized words throughout the conversation, a single conversation unit was fed into the model, enabling unidirectional models to aggregate representations of preceding tokens. Timepoints without word assignments were defined as 0. The resulting concatenated vectors were downsampled to 1 Hz to match the TR. Finally, 12 different stimulus representations (corresponding to the number of GPT-2 layers) consisting of 768 dimensions per conversation were obtained.

In addition, we used two other language models with distinct architectures, namely XLNet^46^ and Llama 2^47^. While both are based on the Transformer, each has its own contextualization mechanisms. XLNet dynamically changes input sequence orders and captures long-term dependencies using the Transformer-XL architecture whereas Llama 2 has high adaptability to various language tasks by learning from large datasets. In the study, like for GPT-2, stimuli transcription was tokenized and input into pretrained models (“xlnet-base-cased,” “meta-llama/Llama-2-7b-hf”) provided by Hugging Face. With XLNet, we obtained 12 different stimulus representations (corresponding to the number of XLNet layers, each with 768 dimensions) for each conversation. For Llama 2, the original features (4,096 dimensions) were reduced to 473 dimensions using principal component analysis, retaining 99% of the information. Additionally, since Llama 2 has 32 layers, we selected 12 layers (1, 4, 7, 9, 12, 15, 18, 21, 24, 26, 29, 32 layers) for analyses.

### 4.8. Encoding model fitting

We constructed encoding models for each of LLM layer and individual. Cortical activity in each voxel was fitted using a finite impulse response model designed to capture the gradual hemodynamic response and its coupling with neural activity^22,80^. The feature matrix **F** [T × 5 N] was modeled by concatenating sets of [T × N] feature matrices with five temporal delays from 2 to 6 s (T = # of samples; N = # of features). The cortical response **R** [T × V] was then modeled by multiplying the feature matrix **F** by the weight matrix **W** [5 N × V] (V = # of voxels):

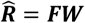

We used an L2-regularized linear regression using the training dataset to obtain the weight matrix **W**. The parameter tunings were conducted in the 10-fold nested cross-validation framework. Training data comprised 2,259 samples and test data 250 samples. The optimal regularization parameter was assessed using seven different regularization parameters from 10 to 10 × 2^6^.

The prediction accuracy was evaluated using the Pearson’s correlation coefficient between predicted and actual signals within the test dataset. The average accuracy was calculated across 10 folds, and the statistical significance (one-sided) computed by comparing the estimated correlations to the null distribution of correlations between two independent Gaussian random vectors of the same length as the test dataset^6,22^. The statistical threshold was set at *p* < 0.05 and corrected for multiple comparisons using the FDR procedure^81^. For subsequent analyses, prediction accuracy for voxels falling below the threshold was considered zero.

### 4.9. Identifying the best-fitting model for each voxel and calculation of model proportions within each ROI

To investigate the relationship between LLM layers and the hierarchy of language representation in individual voxels, we identified the encoding model that demonstrated the highest prediction accuracy from a set of 12 encoding models constructed based on the LLM layers for each participant’s voxels. This was repeated for each LLM (GPT-2, XLNet, and Llama 2) and the identified encoding model was defined as the “best-fit model” for that voxel. The distributions of best-fit models were mapped onto each participant’s brain using pycortex^82^. To determine how each encoding model fits within language-related anatomical ROIs, we further computed the ratio of significant voxels within each ROI for each individual and encoding model.

### 4.10. Correlating prediction accuracy with L2 proficiency in hierarchical GPT-2 layers

For each of the 148 anatomical ROIs and encoding models, we computed the Pearson’s correlation coefficient between EF SET listening scores and the average prediction accuracy of the encoding model within each ROI. Statistical significance (two-tailed) was determined based on the null distribution of correlation coefficients between two independent Gaussian random vectors of the same length as the test dataset. The statistical threshold was set at *p* < 0.05. Corrections for multiple comparisons were applied using FDR^81^, and the number of hypotheses was set to the number of ROIs multiplied by that of models (i.e., 148*12).

### 4.11. Visualization

For data visualization on the cortical maps, we used pycortex^82^ and fsbrain^83^.

## Acknowledgments

We thank MEXT/JSPS KAKENHI (grant numbers JP24H02172 and JP24H01559 for T.N., JP18H05522 and JP24H00619 for S.N.), JST ERATO JPMJER1801, AIP JPMJCR24U2 (for S.N.), and JST FOREST Program (JPMJFR231V for T.N.) for partial financial support of this study. The funders had no role in the study design, data collection and analysis, decision to publish, or preparation of the manuscript.

## Author contributions

**R.K**.: Conceptualization, Data collection, Methodology, Formal analysis, Visualization, Writing– Original draft preparation. **S.N**.: Supervision, Writing–Review & Editing, Funding acquisition. **T.N**.: Supervision, Conceptualization, Writing–Review & Editing.

## Supplementary Information

**Fig. S1.**
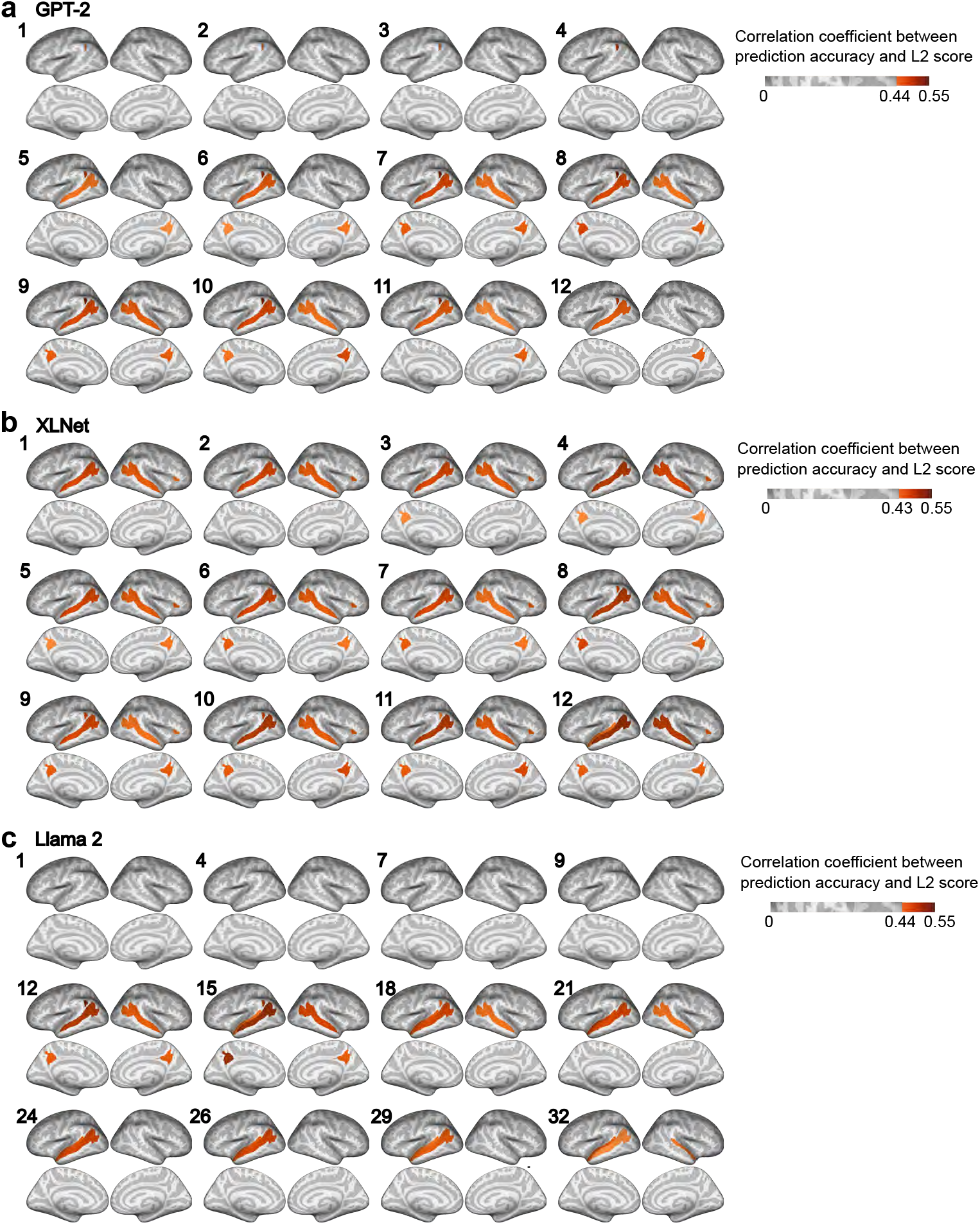
Correlation between prediction accuracy and L2 proficiency. **a** Brain map showing ROIs with significant correlations between L2 proficiency and prediction accuracy (color-coded, *p* < 0.05, FDR corrected). **a** GPT-2. **b** XLNet. **C** Llama 2, 12 representative layers selected at regular intervals from its 32-layer architecture.

